# Imaging single particle profiler to study nanoscale bioparticles using conventional confocal microscopy

**DOI:** 10.1101/2024.06.18.599473

**Authors:** Taras Sych, André Görgens, Loïc Steiner, Gozde Gucluer, Ylva Huge, Farhood Alamdari, Markus Johansson, Firas Aljabery, Amir Sherif, Susanne Gabrielsson, Samir EL Andaloussi, Erdinc Sezgin

## Abstract

Single particle profiler is a unique methodology to study nanoscale bioparticles such as liposomes, lipid nanoparticles, extracellular vesicles and lipoproteins in single particle and high throughput manner. The original version requires the single photon counting modules for data acquisition. Here, we present imaging-based SPP (iSPP) which can be performed by imaging a spot over time in common imaging mode with photomultiplier tubes. We also provide a user-friendly software with graphical user interface to facilitate the application of this technique. We demonstrate that iSPP can be used to decipher lipid-protein interactions, membrane modifications by drugs and the heterogeneity of extracellular vesicles isolated from cells lines and urine of human donors.

## 1. Introduction

High throughput screening of biological particles has wide applications in basic and translational research, and they are commonly used as specific disease biomarkers. For cells, flow cytometry is a widely used technique that allows for analysing tens of thousands of individual cells within minutes, providing information on their type and content. Besides cells, other bioparticles such as lipoproteins^[1]^ (LPs) and extracellular vesicles^[2]^ (EVs) are also commonly used in biomedical research. Quantities of distinct LP types are regularly checked in routine medical examinations to diagnose and prevent metabolic and cardiovascular diseases. Furthermore, it is known that EVs can contain surface proteins designated as disease biomarkers^[3,4]^.

High throughput screening of such biological particles in single particle manner is challenging due to their heterogeneity and small size (10 – 200 nm). There are a few commercially available instruments for this purpose on the market^[5–7]^; however, they are costly and limited in acquisition modalities and analysis. Moreover, they do not perform well with smaller particles. We previously developed a high throughput technique, called single particle profiler (SPP), for profiling nanometre-sized particles based on widely available confocal microscopy^[8]^. We used it for studying artificial liposomes and lipid nano-particles as well as bioparticles obtained from biological material: lipoproteins, extracellular vesicles and virus-like particles. Data for SPP is acquired using confocal microscope; however, it still requires single photon counting detectors that is usually not included in the base models of the confocal microscopes^[9,10]^.

Here, we introduce imaging SPP (iSPP) – single particle profiling performed using standard confocal microscopy, which will increase accessibility of this method and reduce the cost of such measurements. We evaluate the performance of iSPP by addressing several biological questions related to protein-lipid interactions, dynamics of drug effect on membranes and the heterogeneity of biological EVs.

## 2. Results and discussion

The data acquisition for SPP relies on the ability to record photons emitted by fluorescently labelled particles, moving through the observation volume over time. Such photon counting setup is often used for fluorescence correlation spectroscopy (FCS), hence the FCS module was initially employed for data acquisition in SPP ^[8]^. This requirement for specific photon counting detectors and related software adaptation can be a limiting factor since many confocal microscopes designed for imaging (but not for FCS) do not have such detection and acquisition. However, it is not the only mode of a confocal microscope which can yield data as “fluorescent signal over time”. By definition, an image acquired with laser scanning confocal microscopy is a collection of single spots arranged in a grid. The number of such spots is defined by the size of the image and by the spatiotemporal resolution of the microscope. Often, to increase the temporal resolution, the dimensions of the image can be reduced. For example, imaging can be performed in only one plane (no z-axis) or even in a single line (only x-axis) by scanning the same line in one direction over time, which, for example, can be used for diffusion measurements in cellular membranes ^[11]^. Finally, to record fluorescent signal in one spot over time, there is a mode of “spot scan”. For this, the objective is parked in one position and fluorescence intensity is recorded over time (Figure 1a). We utilized this spot imaging for SPP measurements.

**Figure 1.**
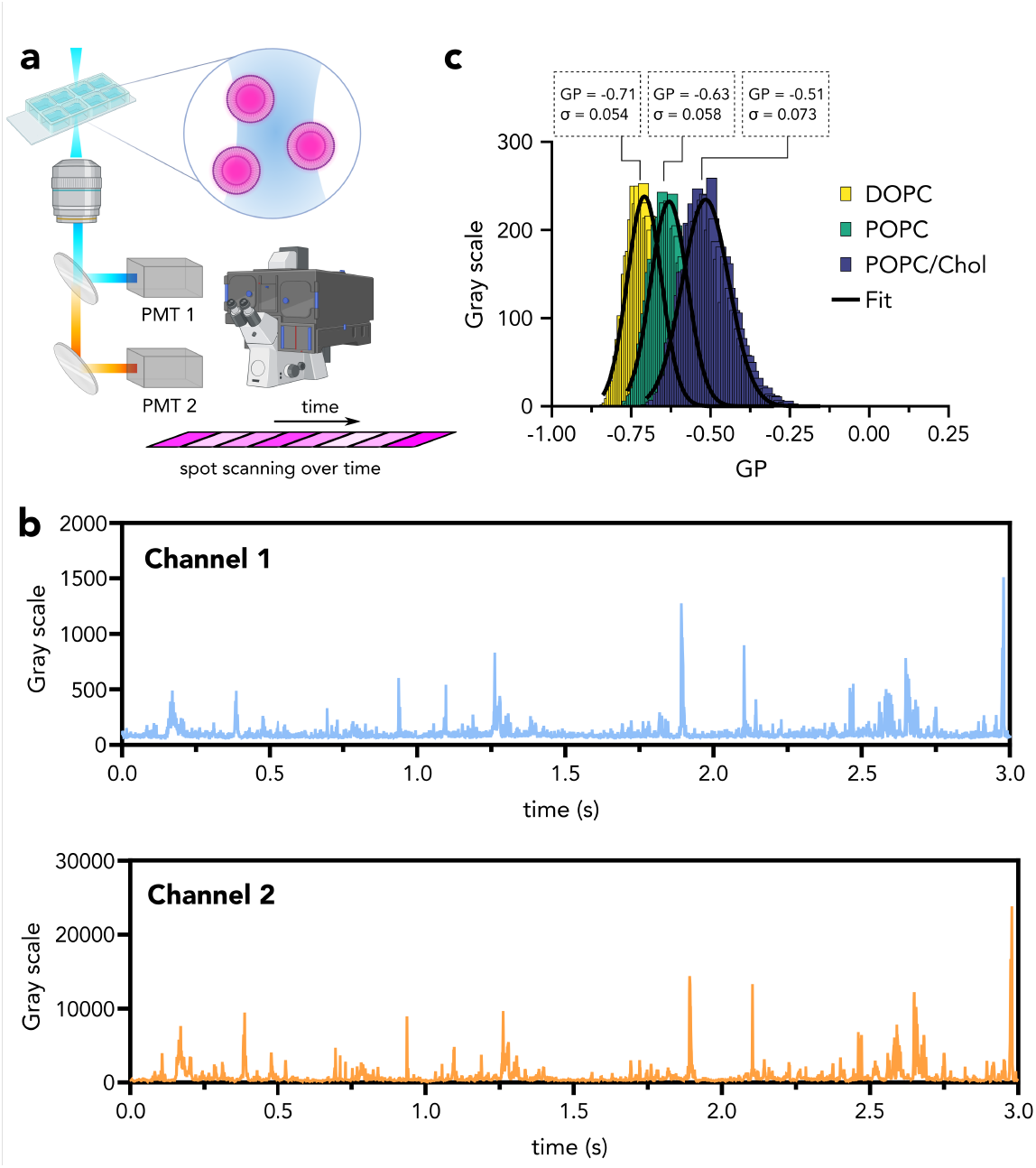
Spot-SPP principle. **a)** The objective is fixed on a spot and PMT detectors record the fluorescence signal over time in multiple channels in imaging mode. Particles are diffusing through the observation spot hence the signal provides intensity fluctuations over time. **b)** Fluorescence fluctuations recorded over time in two channels. **c)** Synthetic liposomes with distinct lipid compositions (pure DOPC, pure POPC and POPC/chol (70/30)) are labeled with environment sensitive probe NR12S and fluorescence intensities for single particles in “green” and “red” channels were recorded. The resulting generalized polarization (GP) histograms for different lipid compositions are presented.

### 2.1. Spot iSPP makes single particle profiling more accessible

The major difference between the “spot scan” and the “single photon mode” is the use of photomultiplier detectors (PMTs, Figure 1a) in “spot scan” vs the photon counting detectors in single photon mode. While measurements with PMTs will make SPP significantly more accessible, there are a few points to consider for reliable data acquisition (link to the detailed video tutorial can be found in Material and Methods section). PMT detector sensitivity can be altered using the voltage gain, hence it is particularly important to keep gain settings the same for different samples that will be compared quantitatively. The same applies for excitation laser power when intensities are directly compared. In “spot scan”, the intensities are recorded in “gray scale” values. Dynamic range here can be increased by increasing the bitrate of the image. As in any confocal images it is important to avoid under- and oversaturation by adjusting laser power, offset and voltage gain of the detector.

Spot iSPP analyzes the fluorescence fluctuation recorded using “spot scan”, i.e., one spot over time. The initial feasibility was evaluated using synthetic liposomes labeled with environmental sensitive dye NR12S ^[12]^. NR12S changes its emission wavelength depending on the fluidity of the membranes. Liposomes doped with NR12A are excellent samples to evaluate the robustness of single particle profiling since liposomes with different lipid compositions have known membrane fluidity which can be used to evaluate the sensitivity of the technique. Moreover, while two emission windows are used for NR12S to calculate the membrane fluidity, only one excitation wavelength is used, which makes the acquisition robust and easy. For this reason, fluorescence intensity traces in two channels were collected where every peak stands for single liposomes passing through the observation volume (Figure 1b). To evaluate membrane fluidity of each liposome, generalized polarization (GP) value was calculated for every peak using the red and green shifted emission wavelengths (see Experimental Section, §4.6 for details) and the GP histograms for different lipid compositions were generated as shown in Figure 1c. Liposomes constituted of pure 1,2-Dioleoyl-sn-glycero-3-phosphocholine (DOPC) show the lowest mean GP, whereas that of 1-palmitoyl-2-oleoyl-sn-glycero-3-phosphocholine and cholesterol (POPC/Chol) is highest. GP distributions are fitted with gaussian distributions, indicating single population for each sample. However, standard deviation (designated as “σ” which is the full width at half maxima) for POPC/Chol is higher than for the other two, displaying higher heterogeneity of membrane fluidity within this sample. This is due to imperfect mixing of lipids in multi-component systems ^[13]^, hence σ in SPP measurements can be used to investigate the sample heterogeneity^[14]^.

Mean GP values follow the same trend as in our previous work using conventional SPP despite different absolute values ^[8]^. This occurs when ratiometric GP measurements are performed at different microscopes, or as in this case on the same device but using imaging mode instead of photon-counting mode. Sensitivities of the detectors are different in different parts of the spectrum, so the absolute GP values might not be preserved. As a result, it is important to use certain “GP standards” in case datasets acquired on different devices will be compared.

### 2.2. Quantification of protein binding to lipid particles with spot iSPP

SPP is not limited to the ratiometric measurements, as done in Figure 1, and can be employed to explore the absolute fluorescence intensities of single particles. A useful application of this type of analysis is the possibility to evaluate the amount of cargo within a vehicle particle (as we previously showed for RNA-loaded LNPs ^[8]^). We also previously demonstrated that SPP can measure binding of different nanobodies to different virus-like particles ^[8]^ or even estimate the number of antibodies bound to a single EV ^[15]^.

To test the performance of Spot iSPP for such molecular quantification, we studied the impact of cholesterol on lipid-protein interactions, namely on binding of carbohydrate-binding proteins (lectins) to their glycosphingolipid receptors. We chose two toxins of the AB^5^ family: Cholera toxin (Ctx) from *V. cholerae* and Shiga toxin (Stx) from *S. Dysenteriae*. Both toxins act similarly: they are taken up by cells and follow the retrograde pathway to trans-Golgi network to block biosynthesis of the proteins in the endoplasmic reticulum. Their cellular uptake is initiated at the plasma membrane by interaction of their B-subunits with corresponding glycosphingolipids (GSLs): globotriaosylceramide (Gb3) for StxB and monosialotetrahexosylganglioside (GM1) for CtxB. Lipid bilayer composition modulates the exposure of carbohydrate moieties to the extracellular space and as a result alters the binding efficiency of toxins to their corresponding receptors. The studies of exact role of cholesterol in toxin-receptor interactions were performed in live cells, by synthetic reconstitution and molecular dynamic simulations. Lingwood et al. ^[16]^ demonstrated that binding of both StxB and CtxB was hindered by cholesterol in erythrocyte plasma membranes as well as in POPC/cholesterol membranes *in silico*. On the other hand, by reconstituting StxB – Gb3 binding in synthetic membranes, Schubert et al. ^[17]^ showed that inclusion of cholesterol in DOPC liposomes promotes StxB binding to Gb3 receptor.

To address this controversy, we reconstituted binding of toxins to their receptors in synthetic liposomes composed of DOPC and 5 mol % of respective GSL with or without 30 mol % of cholesterol (Figure 2a). Liposomes were further supplemented with 0.01 mol % of Fast DiO fluorescent probe to label the membrane while the toxins were labeled with Alexa 647. The traces recorded by Spot iSPP in lipid (Fast DiO) and protein (Alexa 647) channels are shown in Figure 2b. We detected liposomes in lipid channel and quantified the signal in the protein channel for every liposome. The fluorescence signals showed striking effect of cholesterol in protein binding (Supplementary Figure S1). To convert fluorescence signal from proteins into number of proteins per liposome, molecular brightness for both StxB-Alexa647 and CtxB-Alexa647 were calculated by measuring the fluctuations of both proteins in solution (Supplementary Figure S1). Of note, the molecular brightness was assessed by extracting the autocorrelation function from the intensity traces recorded from the proteins in solution also in spot imaging mode, without the photon counting detectors.

**Figure 2.**
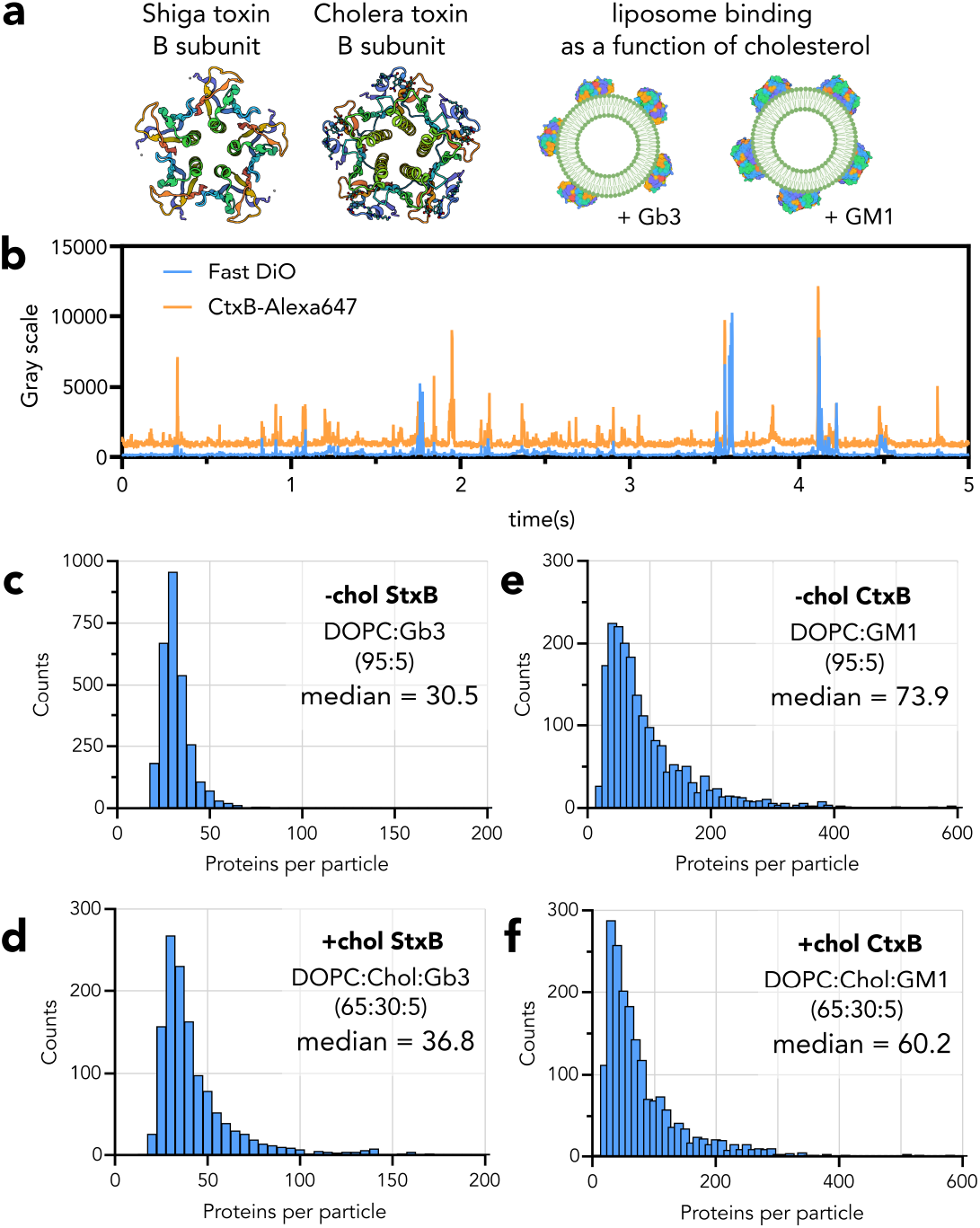
Studying lipid-protein interactions with spot iSPP. **a)** Synthetic liposomes were prepared using the lipid mixtures of DOPC and cholesterol (0 or 30 mol %) with 5 mol % of Glycosphingolipids (Gb3 or GM1). StxB and CtxB were labeled by Alexa 647 and liposomes contained Fast DiO. **b)** Intensity traces in Fast DiO and Alexa 647 channels for CtxB binding experiment to liposomes containing DOPC/GM1 (95/5). **c, d)** Protein per liposome analysis for StxB binding to the liposomes containing DOPC/Gb3 without and with cholesterol, respectively. The histogram is shifted to the right when cholesterol is present which indicates an increase of StxB per liposome. **e, f)** Protein per liposome analysis for CtxB binding to the liposomes containing DOPC/GM1 without and with cholesterol, respectively. The histogram is shifted to the left when cholesterol is present which indicates a decrease of CtxB per liposome.

The histograms of numbers of proteins per liposome are presented in Figure 2c-f. It is evident that for StxB-Gb3 interactions, cholesterol improved the binding of the protein to its receptor (Figure 2c,d), whereas for CtxB, the opposite trend was observed: absence of cholesterol resulted in better binding of CtxB to DOPC liposomes (Figure 2e,f). By analyzing the co-occurrence of lipid and protein signal, we also quantified the percentage of liposomes bound by proteins for every condition (Supplementary Table S1). For StxB-Gb3, 29% of cholesterol-positive and 20% of cholesterol-negative liposomes were bound by proteins. For CtxB-GM1, 52% of cholesterol-positive liposomes and 60% of cholesterol-negative liposomes were bound by CtxB. These results are fully in line with previous studies showing cholesterol hindering the CtxB binding. For StxB, however, cholesterol showed the improvement of StxB binding, which is in line with Schubert et al. ^[17]^, but contradicts the Lingwood et al. ^[16]^ which could be due to the different base lipid used (DOPC vs. POPC). Importantly, it was not possible to remove free proteins from the solution, hence they create background in the Alexa 647 channel (Figure 2b). This background value can mask detection of protein binding to liposomes, in case the number of proteins per liposome is closer to the background which can change the absolute percentage.

### 2.3. Dissecting the dynamics of membrane remodeling upon drug interactions

Spot iSPP can profile several thousands of particles in a few minutes. Moreover, within short timeframes (≈2 min), it is possible to profile enough particles to resolve multiple subpopulations within the data distribution. This opens the possibility to perform several consecutive short-time acquisitions and as a result to explore the dynamic processes, such as drug interaction with lipid bilayers. One widely used drug for manipulation of membrane composition is methyl-ß-cyclodextrin (MBCD). Cholesterol encapsulation by MBCD is widely used to remove cholesterol from native and artificial lipid bilayers, which is a standard procedure when the role of cholesterol in various biological processes is in question ^[16,18–20]^. Interestingly, it is also possible to use MBCD as an intermediary to deliver and distribute cholesterol homogeneously among artificial liposomes ^[14]^. MBCD interaction with membranes is complex especially when there are multiple different lipid environments present. To test which membrane environments MBCD preferentially interacts with, we prepared synthetic liposomes (Figure 3a) of POPC/Chol (70/30) and DPPC/Chol (70/30) and labeled them using NR12S. The mixture of these liposomes at 1:1 ratio was profiled and resulted in the GP histogram displayed in Figure 3b. This histogram consists of two distinct populations: the first population centered at −0.57 represents POPC/Chol liposomes and the second population at 0.45 which is DPPC/Chol liposomes. When MBCD (2 mM) is applied to POPC/Chol or DPPC/Chol separately, in both cases it extracts some cholesterol, and the GP distribution shifts towards smaller GP values (Supplementary Figure S2). MBCD also extracts some NR12S, but it is unlikely that signal from NR12S extracted by MBCD can contaminate results, because the GP of NR12S incorporated in MBCD is extremely low (around −0.9, Supplementary Figure S2) and such population, if significant, would be clearly visible as a separate population.

**Figure 3.**
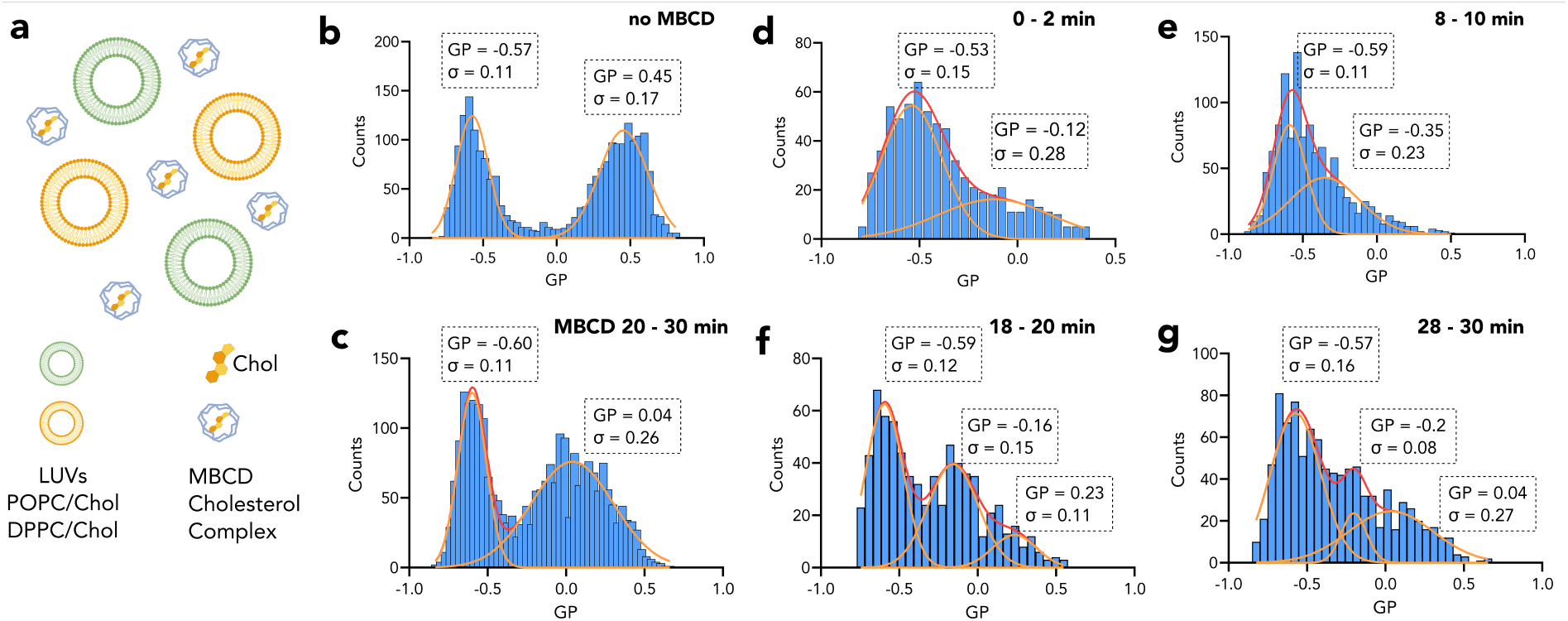
Cholesterol extraction from liposomes by MBCD. **a)** Liposomes of POPC/Chol (70/30) and DPPC/Chol (70/30) were labeled with environmental sensitive probe NR12S and mixed. **b)** GP histogram of POPC/Chol and DPPC/Chol liposomes mixed in the well plate at ratio 1:1. **c)** GP histogram of POPC/Chol and DPPC/Chol mixture recorded between 20-30 min after MBCD application. **d-g)** GP histogram of POPC/Chol and DPPC/Chol mixture recorded between **d)** 0-2 min, **e)** 8-10 min **f)** 18-20 min **g)** 28-30 min after MBCD application.

However, interestingly, when MBCD (2 mM, 30 min) is applied to the mixture of POPC/Chol and DPPC/Chol, the population of POPC/Chol remains unchanged, whereas DPPC/Chol population shifts towards smaller GP values significantly (Figure 3c). It seems that MBCD prefers to extract cholesterol from DPPC/Chol liposomes rather than from POPC/Chol liposomes. Shaw et al. ^[21]^ studied similar problem by assessment of the chemical potential differences between POPC/Chol and DPPC/Chol at different cholesterol content. According to their study, however, chemical potential of POPC/Chol was higher compared to chemical potential of DPPC/Chol at cholesterol concentration of 30 mol %. This means that MBCD would more likely extract cholesterol from POPC/Chol first. DPPC/Chol has higher chemical potential than POPC/Chol at cholesterol concentration of at least 40-50 mol %. This necessitates the time-lapse measurements to observe the time evolution of cholesterol extraction from different populations. Since we have access to various timepoints after MBCD application to the liposome mixture, we have insights on the dynamics of cholesterol extraction by MBCD. For this, we have recorded the intensity traces for 30 minutes continuously after 2 mM MBCD application to the liposome mixture, GP histogram of which before MBCD application is displayed in Figure 3b. After the acquisition was completed, the GP histograms were constructed for the shorter time traces of 2-min. These histograms represent transient states appearing and disappearing before it reaches an equilibrium. Immediately after MBCD application (0-2 min), cholesterol is being extracted from DPPC/Chol liposomes and population of large GP values shifts left (Figure 3d). Moreover, at 8-10 min timepoint, the high GP population is practically absent (Figure 3e). Later, however, the higher GP population reappears. At 20-22 min (Figure 3f) and at 28-30 min (Figure 3g) timepoints, there are more than two populations: lowest GP population still aligns well with POPC/Chol (GP ≈ −0.57), whereas the other two populations indicate large heterogeneity. Again, the traces acquired between 20 - 30 minutes (Figure 3c) have quite broad GP distribution. This population can consist of multiple minor sub-populations that are better pronounced in the transient histograms in Figure 3f and 3g. Of note, the POPC/Chol population changes little to none during 30 min after MBCD application.

This data shows that drug interactions with membrane can be a complex phenomenon with multi-step dynamics and intermediate steps. Specifically, MBCD interaction with membranes occurs as a function of the chemical potential of the membrane systems and MBCD itself. Therefore, during the cholesterol removal by MBCD, there could be many intermediate chemical potential systems that play a role in drug action.

### 2.4. Biological particles from cultured cells and body fluids

Biological nanosized particles (such as lipoproteins and extracellular vesicles) are widely used in basic and translational research and as biomarkers of diseases ^[4,22]^. Their small size and heterogeneity make it challenging to study them in single particle and high throughput manner. Therefore, we aimed to use spot iSPP for studying nanoscale biological particles to profile their content and biophysical properties. To this end, we studied membrane fluidity of EVs. EVs are small particles secreted by cells and used for cellular communication ^[23,24]^. Recently, they are also designed for therapeutic purposes ^[15,25,26]^. To be able to use them in biomedical research, it is crucial to shed light on their content heterogeneity. We extracted EVs from different cultured cells (Figure 4a-c) and labelled them with NR12S. The membrane fluidity analysis with iSPP resulted in one component GP distributions (Figure 4b) for all but with different fluidities. EVs isolated from HUVEC endothelial and HUAEC epithelial cell lines exhibited the most fluid membranes, whereas the GP of MSC-EVs was slightly higher. Fibroblast-derived EVs showed similar GP as T-lymphocytic EVs from Jurkat cells (the only suspension cell line in this collection), which were the least fluid EVs. This shows that depending on the cell source, the membrane content of the EVs varies dramatically. Moreover, biophysical heterogeneity should be considered for the efficacy of EV delivery and uptake by target cells.

**Figure 4.**
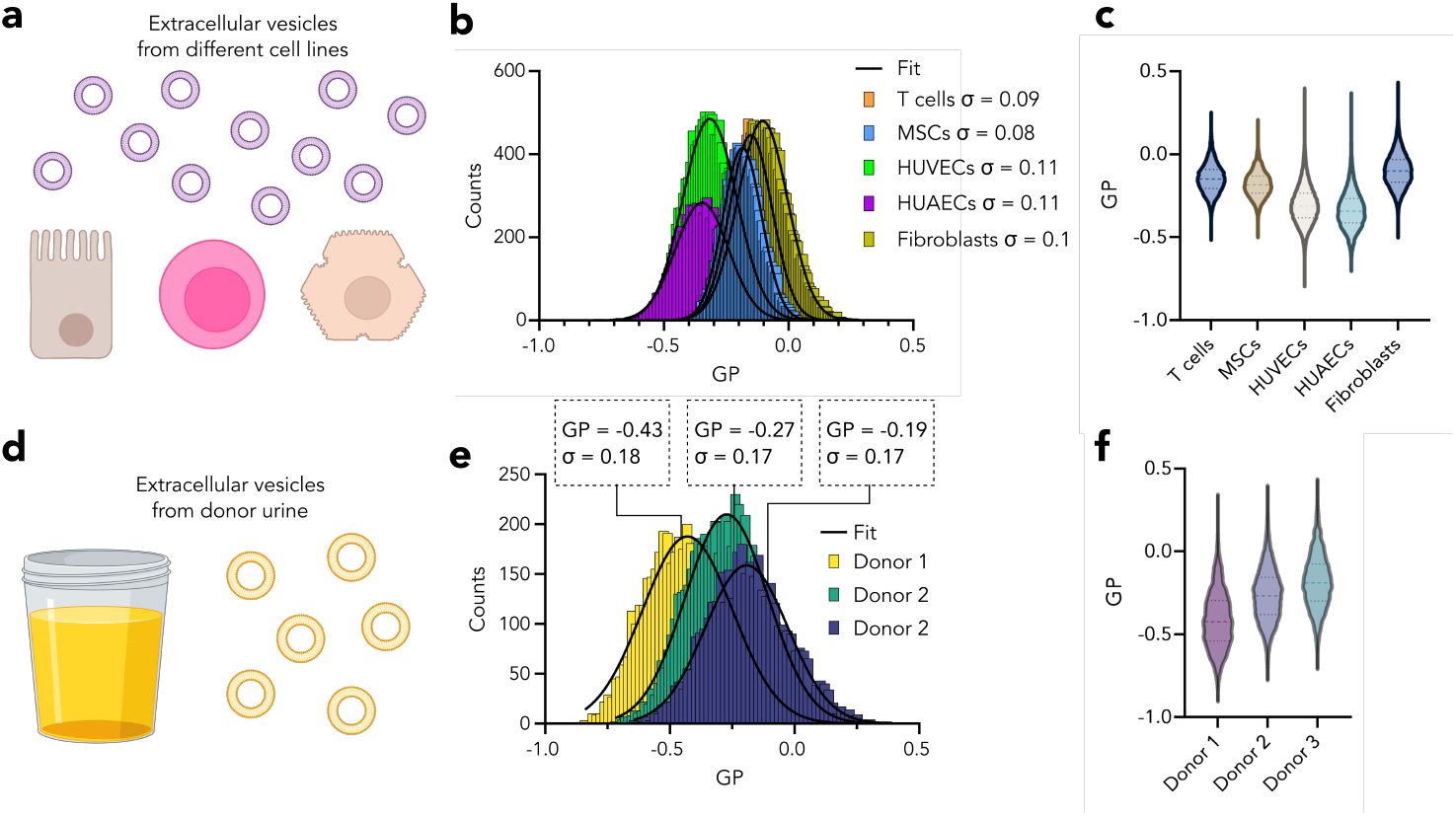
Membrane fluidity of extracellular vesicles. **a)** EVs were isolated from T cells, MSCs, HUVECs, HUAECs and fibroblasts, labeled with the environment sensitive probe NR12S and profiled using spot iSPP. **b)** GP histograms of EVs isolated from various cells in culture. **c)** GP violin plot of EVs in panel b. **d)** EVs were isolated from urine of three donors, labeled with NR12S and profiled using spot iSPP. **e)** GP histograms of EVs isolated from donor urine. **f)** GP violin plot of EVs in panel e.

To show that iSPP can also be used in more physiological setting for nanoscale bioparticle characterization, we used extracellular vesicles extracted from human donor urine. EVs were isolated from three different donors and showed significant differences in their fluidity. This shows that iSPP can be used to study properties and heterogeneity of nanoscale bioparticles from body fluids and can successfully show individual differences which can be a new avenue for biomarker discovery.

## 3. Conclusion

Nanoscale bioparticles are important biomarkers for diseases and crucial tools for biomedicine. Therefore, their content, heterogeneity and properties have significant implications for understanding physiology and diseases as well as using them as biomedical tools. Methodologies to study these particles were limited; however, there are several initiatives to overcome this technological bottleneck. SPP is one of these techniques we developed recently to make single particle measurements possible on commercially available microscopes in a high throughput and single particle manner.

In this work, we develop a new version of SPP – Spot imaging SPP (iSPP), which is more accessible for users with less advanced microscopy experience as well as for users with limited access to advanced confocal microscopes equipped with photon counting detectors. Spot iSPP was validated here with on artificial and native lipid particles for two-color profiling as well as for biophysical characterization. It is available for microscopes with photo multiplying detectors and will assist in single particle analysis of bioparticles in health and disease, for lipid and protein profiling as well as for deciphering of biophysical disease biomarkers.

## 4. Experimental Section/Methods

### 4.1. Preparation of LUVs

1,2-Dioleoyl-sn-glycero-3-phosphocholine (DOPC), 1-palmitoyl-2-oleoyl-sn-glycero-3-phosphocholine (POPC), Dipalmitoylphosphatidylcholine (DPPC) and cholesterol (Chol) were from Avanti polar lipids. Glycosphingolipid globtriaosylceramide (Gb3) was from Larodan, monosialotetrahexosylganglioside was from Sigma. Fluorescent lipids and lipid-like probes we used: FastDiO (ThermoFisher), NR12S (Bio-Techne). Lipid mixtures in chloroform at 0.25 mg/ml were prepared: pure DOPC, pure POPC, POPC/ Chol (70/30), DPPC/ Chol (70/30). DOPC/Chol/Gb3 (65/30/5), DOPC/Gb3 (95/5), DOPC/Chol/GM1 (65/30/5), DOPC/GM1 (95/5). Mixture including GM1 or Gb3 were further supplemented with 0.01 mol % of FastDIO. Mixtures were dried under the flow of nitrogen, rehydrated with buffer (150 mM NaCl, 10 mM Hepes, 2 mM CaCl_2_) and vortexed harshly to form multilamellar vesicles. Then, the suspension of MLVs was sonicated at power 3, duty cycle 40% for 10 mins using Branson Sonifier 250. LUVs were stored at 4 degrees under nitrogen. For membrane fluidity studies, the solutions of LUVs (0.25 mg/ml) were incubated with 1 µM Nile Red 12 S (NR12S, stock concentration of 10 µM in DMSO) directly before profiling.

### 4.2. Lectin labelling and application

The pentameric B-subunit of Shiga toxin (StxB) and the pentameric B-subunit of Cholera toxin were purchased from Sigma Aldrich. Lectins were labeled with Alexa-647 (NHS ester) from Thermofisher. Briefly, 1 μL of a 10 mg/mL solution of the amino-reactive probe dissolved in DMSO was added to 100 μL of a 0.5 mg/mL protein solution in PBS (-/-, Gibco) supplemented with 100 μM NaHCO3, pH 8.5. The mixture was incubated for 1 h at room temperature under continuous shaking. The labeled lectins were purified using Zeba Spin Desalting Columns (0.5 mL, MW cut-off: 7.0 kDa) from ThermoFisher Scientific.

For measurements of StxB and CtxB in solution they were brought to the concentration of 100 nM in PBS. For binding studies 10 μl of LUVs that contained appropriate GSL receptor were incubated with 1 μM of lectin for 15 minutes before profiling with Spot SPP.

### 4.3. MBCD application

Methyl-ß-cyclodextrin (MBCD) was from Sigma. MBCD was freshly dissolved with ultrapure water at a concentration of 40 mM. For control experiments, LUVs (POPC/Chol or DPPC/Chol) were incubated for 30 min with 2 mM of MBCD and diluted 10 times for profiling with Spot SPP. For control experiment on fluorescent dye capture by MBCD, 2 mM of MBCD was mixed with 1 μM solution of NR12S, incubated for 30 min and diluted 10 times for Spot SPP profiling. For the cholesterol extraction dynamics study, POPC/Chol and DPPC/Chol (1:1) mixture was added to the profiling well plate and the initial 10 min measurement was performed. Later, 2 mM of MBCD was added to exactly the same well, and Spot-SPP profiling was started immediately after MBCD addition. Traces were recorded for 30 minutes and later split into shorter traces in order to analyze the time series.

### 4.4. Cell culture and isolation of extracellular vesicles

EVs were prepared from Jurkat T-lymphocyic suspension cells, CB-MSCs, HUVEC, HUAEC, and fibroblasts. Cell lines were cultured in the following media: Jurkat T cells were cultured in RPMI1640 medium. CBMSCs (MSCs; ATCC, PCS-500-010, Umbilical Cord-Derived Mesenchymal Stem Cells, Normal, Human) were cultured in MEM-α modification medium (containing L-glutamine; ThermoFisher Scientific) supplemented with 5 ng/ml of bFGF (Sigma, F0291). Immortalized human umbilical vein endothelial cells (HUVEC/TERT2; ATCC #CRL-4053) were cultured in Vascular Cell Basal Medium (ATCC #PCS-100-030) supplemented with endothelial cell growth Kit-VEGF (ATCC #PCS-110-041). HUAEC (CI-huAEC; Inscreenex, human airway epithelial cells) were cultured in huAEC media (INS-ME-1013-500ml; Inscreenex) supplemented with 1% sodium pyruvate (ThermoFisher). BJ-5ta fibroblast cells (ATCC CRL-4001) were cultured with 4:1 mixture of Dulbecco’s modified Eagle’s medium (containing 4 mM L-glutamine, 4.5 g/L glucose and 1.5 g/L sodium bicarbonate) and Medium 199 (0.01 mg/ml Hygromycin B/10687010, Thermo Fisher) supplemented with 10% FBS (ThermoFisher). Unless indicated otherwise, all cells were supplemented with 10% FBS (Invitrogen) and 1X Antibiotic-Antimycotic (Anti-Anti) (ThermoFisher Scientific). All cell lines were grown at 37°C, 5% CO_2_ in a humidified atmosphere and regularly tested for the presence of mycoplasma. For EV harvesting, cell culture-derived conditioned media (CM) was changed to OptiMem (Invitrogen) 48 h before harvest of conditioned media as described before^[27]^. Unless indicated otherwise, all conditioned media samples were directly subjected to a low-speed centrifugation step at 500 xg for 5 min followed by a 2,000 xg spin for 10 min to remove larger particles and cell debris. Pre-cleared cell culture supernatant was subsequently filtered through 0.22 µm bottle top vacuum filters (Corning, cellulose acetate, low protein binding) to remove any larger particles. EVs were first prepared by tangential flow filtration (TFF) by using a KR2i TFF system (SpectrumLabs) equipped with modified polyethersulfone (mPES) hollow fiber filters with 300 kDa membrane pore size (MidiKros, 370 cm2 surface area, SpectrumLabs) at a flow rate of 100 mL/min (transmembrane pressure at 3.0 psi and shear rate at 3700 sec−1) as described previously ^[28]^. Amicon Ultra-0.5 10 kDa MWCO spin-filters (Millipore) were used to concentrate the sample to a final volume of 100 µL. The sample was then loaded on a qEV column (Izon Science) and the EV fractions were collected according to the manufacturer’s instructions. Final EV samples were stored at −80°C in PBS-HAT [PBS supplemented with HEPES, human serum albumin and D-(+)-Trehalose dihydrate] buffer until usage^[29]^.

### 4.5. Urine collection and extracellular vesicles isolation

Three patients with non-muscle invasive urothelial urinary bladder cancer, staged TaG1, TaG2 and T1G3 respectively, were prospectively recruited between 2016 and 2018 and urine was obtained prior to the primary transurethral resection of the bladder tumor (TURBT). The three patients were 69, 61 and 78 years of age: two females and one male. All samples were shipped and processed fresh the day of surgery. All experimental protocols were approved by the Regional Ethical Review Board in Stockholm (original no.: 2007/71-31), and all patients were above 18 and gave written and oral informed consent. Urine samples were sequentially centrifuged at 300 xg for 10 min, 3000 xg for 30 min, 10,000 xg (Ti45 rotor, Beckman Coulter, tube average) for 30 min and filtered through a 0.22µm filter. The urine was then up concentrated by tangential flow filtration (100kDa cut-off, VivaFlow). After storage at −80ºC, extracellular vesicles were purified by ultracentrifugation at 100,000 xg (Ti45 rotor, Beckman Coulter, tube average) for 2h, and the pellet was resuspended in a small volume of PBS and stored at −80ºC until further analysis.

### 4.6. Single particle profiling measurements and analysis

Single Particle Profiling was performed using the setup for confocal acquisition („spot scan”) at a Zeiss LSM 780 microscope. A 488 nm argon ion laser was used for FastDiO and NR12S excitation, whereas a 633 nm He−Ne laser was used for Alexa 647. A 40×1.2 NA water immersion objective was used to focus the light. The laser power was set to 1% of the total laser power that corresponds to 20 μW. The fluorescence emission was detected by GaAsP spectral detector in integration mode, with HV Gain of 900 – 1100.The emission detection windows were set as 490 – 560 for FastDiO and 650 – 700 for Alexa647. Emission from NR12S was recorded simultaneously in both channels. Intensity traces were recorded between10 and 30 minutes and later split if required. The detailed description and guide on data acquisition is available as video tutorial here: https://www.youtube.com/watch?v=IicPvjPySDY&list=PLvnnxg3kwLpWX5M-e-14hhWQVPsPEQ5jB.

Traces and curves were then analysed using “Py Profiler”, our in-house python program using the python packages: tkinter (v. 8.6.10), matplotlib (v. 3.3.4), lmfit (v. 1.0.2), ttkwidgets (v. 0.10.0), scipy (v. 1.6.2), seaborn (v. 0.11.1), pandas(v. 1.2.4). The source code as well as the standalone distributions for Windows and Mac are available at the Github: https://github.com/taras-sych/Single-particle-profiler/tree/Release-v-3.0.

### Peak analysis

Briefly, individual peaks from the traces were identified and intensities for these individual peaks were extracted with further calculation of generalized polarization (GP) if applicable. GP was calculated using the formula:

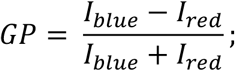

where *I*_*blue*_ – fluorescence intensity in short wavelength („blue”) region of the emission spectrum and *I*_*red*_ – fluorescence intensity in long wavelength („red”) region of the emission spectrum.

The description and guide to the program is available as a video tutorial. The link to the video is at https://www.youtube.com/watch?v=p5afSZSkbfE.

### Diffusion analysis

was also performed for quantification of brightness of proteins in solution using Py Profiler. Curves were fitted with the following three-dimensional diffusion:

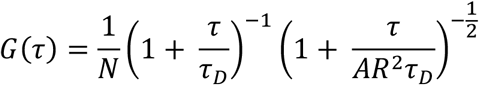

where *N* represents the number of fluorescent species within the beam’s focal volume. Next, molecular brigtness was quantified by dividing the mean value of the trace intensity by *N*.

## Supporting Information

Supporting Information is available from the Wiley Online Library.

## Acknowledgements

We thank Sarah Veatch for the discussion. This work is supported by Swedish Research Council Starting Grant (grant no. 2020-02682), Wellcome Leap’s Dynamic Resilience Program (jointly funded by Temasek Trust), SciLifeLab National COVID-19 Research Program financed by the Knut and Alice Wallenberg Foundation, Cancer Research KI, Human Frontier Science Program (RGP0025/2022) and Longevity Impetus Grant from Norn Group, Hevolution Foundation and Rosenkranz Foundation. We are grateful to Swedish Medical Research Council (VR 2022-01170), The Swedish Cancer Foundation (22-2102 Pj), Cancer Research Foundations of Radiumhemmet (131082 and 181103). We thank the SciLifeLab Advanced Light Microscopy facility and National Microscopy Infrastructure (VR-RFI 2016-00968) for their support on imaging. AG is an International Society for Advancement of Cytometry (ISAC) Marylou Ingram Scholar 2019-2024 and supported by Karolinska Institutet Network Medicine Alliance collaboration grants.

Received: ((will be filled in by the editorial staff))

Revised: ((will be filled in by the editorial staff))

Published online: ((will be filled in by the editorial staff))

## Notes

### Competing Interest Statement

The authors have declared no competing interest.

