## Supplementary Figures for "Imaging single particle profiler to study nanoscale bioparticles using conventional confocal microscopy"

### Supporting Information

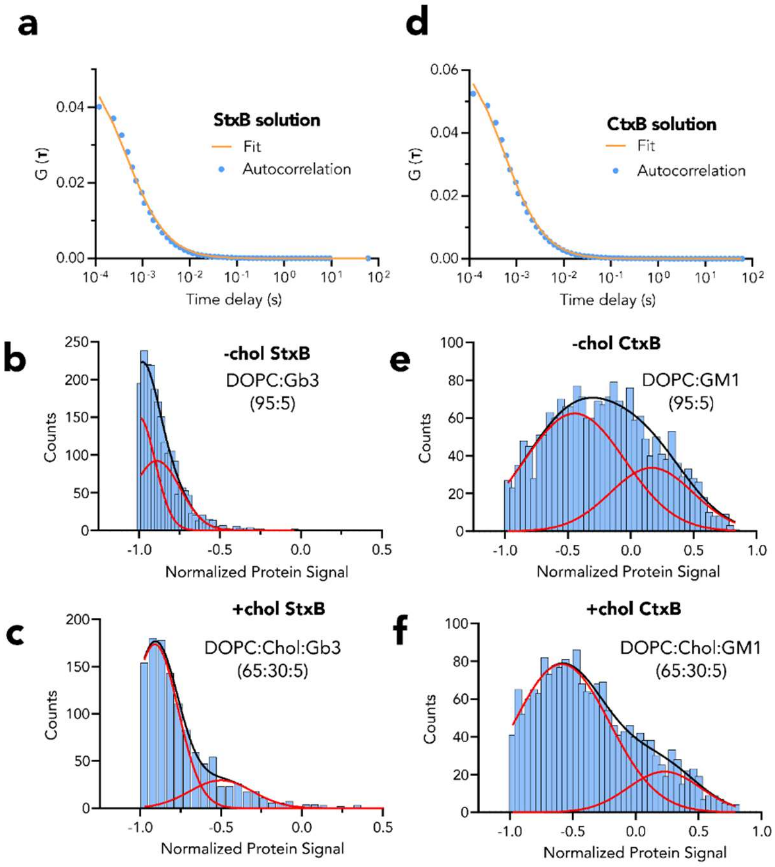


**Figure S1.** Lectin binding to LUVs that contain glycosphingolipid receptors. a) Autocorrelation function ant its fit obtained from fluorescence fluctuations from StxB in solution; b) Autocorrelation function ant its fit obtained from fluorescence fluctuations from CtxB in solution; b-f) ratiometric histograms of protein binding to LUVs with or without cholesterol. Ratio for every particle was calculated by the formula:

$$Ratio = \frac{I_{protein} - I_{protein\ backgound} - I_{lipid}}{I_{protein} - I_{protein\ backgound} + I_{lipid}}$$

**Table S1:** Number of liposomes bound by lectins

| Lipid composition | Lectin | Total number of liposomes | Number of liposomes, bound by lectin | Percentage of liposomes, bound by lectin |
|---|---|---|---|---|
| DOPC/Chol/Gb3 | StxB | 1305 | 374 | 29% |
| DOPC/Gb3 | StxB | 2897 | 585 | 20% |
| DOPC/Chol/GM1 | CtxB | 1954 | 1019 | 52% |
| DOPC/GM1 | CtxB | 2034 | 1222 | 60% |

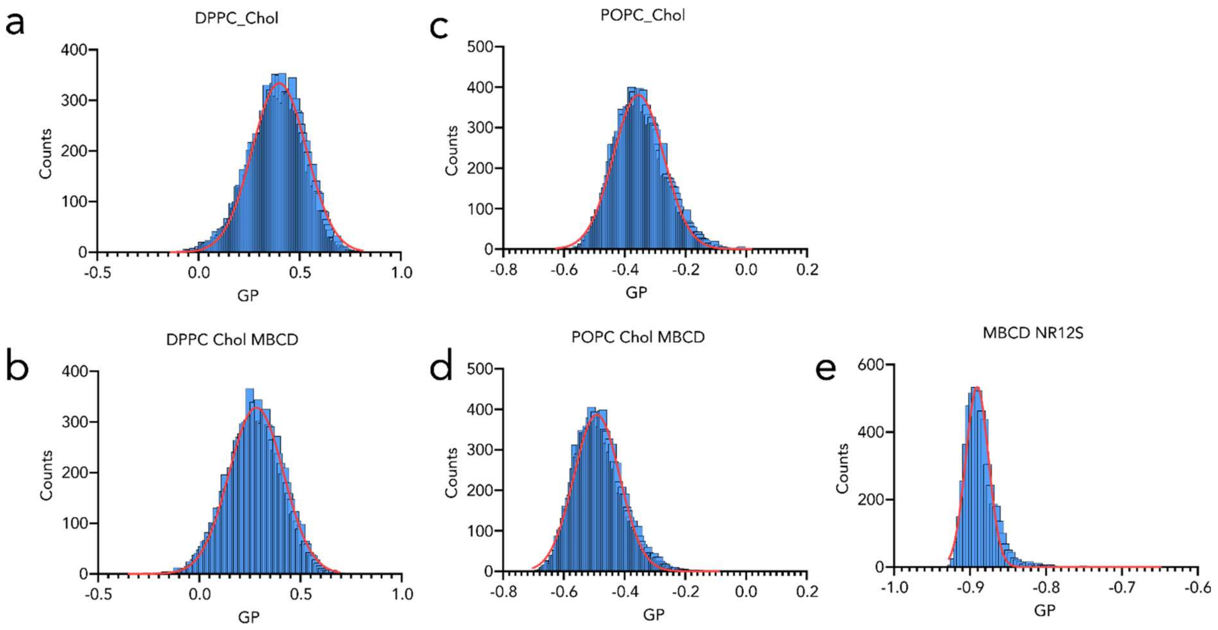


**Figure S2.** MBCD impact on LUVs. a) GP histogram of DPPC/Chol LUVs before 2 mM MBCD application; b) GP histogram of DPPC/Chol LUVs after 2 mM MBCD application; c) GP histogram of POPC/Chol LUVs before 2 mM MBCD application; d) GP histogram of POPC/Chol LUVs after 2 mM MBCD application; e) GP histogram of the solution of NR12S incubated with MBCD.
